# RNA-Rocket: An RNA-Seq Analysis Resource for Infectious Disease Research

**DOI:** 10.1101/007963

**Authors:** Andrew S. Warren, Cristina Aurrecoechea, Brian Brunk, Prerak Desai, Scott Emrich, Gloria I. Giraldo-Calderón, Omar Harb, Deborah Hix, Daniel Lawson, Dustin Machi, Chunhong Mao, Michael McClelland, Eric Nordberg, Maulik Shukla, Leslie B. Vosshall, Alice R. Wattam, Rebecca Will, Hyun Seung Yoo, Bruno Sobral

## Abstract

**Motivation:** RNA-Seq is a method for profiling transcription using high-throughput sequencing and is an important component of many research projects that wish to study transcript isoforms, condition specific expression, and transcriptional structure. The methods, tools, and technologies employed to perform RNA-Seq analysis continue to change, creating a bioinformatics challenge for researchers who wish to exploit these data. Resources that bring together genomic data, analysis tools, educational material, and computational infrastructure can minimize the overhead required of life science researchers.

**Results:** RNA-Rocket is a free service that provides access to RNA-Seq and ChIP-Seq analysis tools for studying infectious diseases. The site makes available thousands of pre-indexed genomes, their annotations, and the ability to stream results to the bioinformatics resources VectorBase, EuPathDB, and PATRIC. The site also provides a combination of experimental data and metadata, examples of pre-computed analysis, step-by-step guides, and a user interface designed to enable both novice and experienced users of RNA-Seq data.

Availability: RNA-Rocket can be found at rnaseq.pathogenportal.org Source code for this project can be found at github.com/cidvbi/PathogenPortal

## 1 INTRODUCTION

Transcriptomic analysis using high-throughput sequencing continues to increase in popularity due to low sequencing costs, its sensitivity, reproducibility, and its ability to sample the entire transcriptome (Wang, Gerstein, & Snyder, 2009). As an active area of research there are many variations of RNA-Seq protocols, data types, and tools that continue to evolve. As a result there is no “one size fits all” solution for doing RNA-Seq analysis. The range of considerations facing life scientists who want to leverage this technology can demand significant investment of time and resources. For this reason we have created RNA-Rocket, an RNA-Seq analysis service that enables infectious disease research for prokaryotic and eukaryotic pathogens as well as vectors, and host genomes.

RNA-Rocket is built on Galaxy (Blankenberg et al., 2001; Giardine et al., 2005; Goecks, Nekrutenko, & Taylor, 2010), with modifications to help simplify the process for routine use and provide a guided user experience. RNA-Rocket integrates data from the PATRIC, EuPathDB, and VectorBase Bioinformatics Resource Centers (BRCs) and is provided by Pathogen Portal (pathogenportal.org), a resource linking all BRCs funded by the National Institute of Allergy and Infectious Diseases (NIAID).

The RNA-Rocket service leverages multiple open source software tools to provide a free resource where users can upload their RNA-Seq data, align them against a genome, and generate quantitative transcript profiles. This service also provides streaming of alignment and annotation results back to the appropriate BRC so that users can view results in the relevant BRC, using annotations and tools provided in support of transcriptomic analysis.

The **Pat**hosystems **R**esource **I**ntegration **C**enter (PATRIC) is the all-bacteria Bioinformatics Resource Center (patricbrc.org) (Wattam et al., 2013). PATRIC provides researchers with an online resource that stores and integrates a variety of data types (e.g. genomics, transcriptomics, protein-protein interactions, 3-D protein structures and sequence typing data) along with any associated metadata. Data types are summarized for individual genomes and across taxonomic levels, and also for individual genes. All genomes in PATRIC, currently more than 21,000, are consistently annotated using Rapid Annotations using Subsystems Technology, RAST (Overbeek et al., 2013). Comparisons of PATRIC and NCBI (Pruitt, Tatusova, Klimke, & Maglott, 2009) annotations are available. PATRIC provides a variety of ways for researchers to find data of interest and a private workspace where they can store both genomic and gene associations, and their own private data, including RNA-Seq data downloaded from RNA-Rocket. Both private and public data can be analyzed together using a suite of tools to perform comparative genomic or transcriptomic analysis.

The eukaryotic pathogen databases (EuPathDB: http://eupathdb.org) provide access to a variety of data types from important human and veterinary parasites such as *Plasmodium* (malaria), *Cryptosporidium* (cryptosporidiosis) and kinetoplastida (i.e., *Trypanosoma brucei* and *Leishmania* species.) (Aurrecoechea et al., 2013). A graphical search system enables users to interrogate underlying data such as genomic sequences, annotation, functional genomic data (i.e., RNA-Seq and proteomic) and clinical data (Fischer et al., 2011). Users of EuPathDB can construct complex search strategies such as asking for genes that contain predicted secretory signal peptides that are expressed during a specific developmental stage, that are absent from mammals and contain a large number of polymorphisms. Results of searches may be viewed or downloaded. In addition, users may upload their own data such as RNA or DNA sequence coverage plots into a genome browser (GMOD), where they can view their uploaded data in the context of their genome of interest with any data already available in EuPathDB (Stein et al., 2002). Data generated through RNA-Rocket are conveniently linked to the genome browser in EuPathDB sites for rapid visualization.

VectorBase (www.vectorbase.org) is a bioinformatics resource for invertebrate vectors of human parasites and pathogens (Megy et al., 2012). It currently hosts the genomes of 35 organisms including mosquitoes (20 of which are *Anopheles* species), tsetse flies, ticks, lice, kissing bugs, and sandflies. VectorBase has also recently incorporated *Biomphalaria glabrata*, the snail intermediate host of schistosomiasis and the house fly, *Musca domestica*. Hosted data range from genomic features and expression data to population genetics and ontologies. Updates to VectorBase (or releases) occur every two months; both public and private data can be queried or analyzed with a variety of available tools including BLAST and Expression and Population Biology browsers (Megy et al., 2012). The overarching goal of VectorBase resource is to support “-omics” aided efforts to improve or develop new vector control strategies.

## 2 METHODS

### 2.1 Software and tools

RNA-Rocket takes advantage of many different open-source projects to enable users to upload and analyze their own data. We use the Galaxy system to consolidate and provide the tools and services necessary to process high-throughput sequencing data. The use of Galaxy has many benefits: showing provenance information for data creation, including the tools and parameters used to process data; support for batch analysis for multiple samples; providing a mechanism for results sharing across research groups and publishing for external references such as presentations or publications; and its integration of tools and projects in the larger bioinformatics community.

Before users can run analysis on the RNA-Rocket site they must first upload their data in FASTQ format. Using standard Galaxy interfaces RNA-Rocket supports upload via URL, FTP, HTTP, and direct transfer via the European Nucleotide Archive (Rasko Leinonen et al., 2011), which can be searched using ENA, SRA, GEO and ArrayExpress identifiers. To enable basic RNA-Seq processing, RNA-Rocket provides users with a set of predetermined Galaxy workflows configured to use existing BRC genomes and annotations. The primary workflow for RNA-Seq analysis aligns short read data to a reference genome using Bowtie2 (Langmead & Salzberg, 2012) or TopHat2 (Kim et al., 2013), assembles transcripts using Cufflinks, and generates coverage bed-Graph and BigWig files using BEDTools (Quinlan & Hall, 2010) and UCSC tools (Kuhn, Haussler, & Kent, 2013) respectively. This workflow generates BAM files and tab-delimited output, which can be used to determine transcript structure and the level of expression in the target organism. The site also provides the ability to conduct differential expression analysis using Cuffdiff, part of the Cufflinks suite, and the ability to visualize data generated using CummeRbund (Trapnell et al., 2012). When users submit their jobs to RNA-Rocket, they are queued and run on a first-come, first-served basis on a compute cluster using modern, high-density, computer architecture.

The site features an interactive concept diagram, which highlights the appropriate processing step(s), based on the concept that the user is interested in. Clicking on a concept diagram component gives information about the corresponding processing steps that fulfill the component (Figure 1). To assist users, the site provides a ‘Launch Pad’ menu system that breaks down the context and rationale for executing a particular step, the input required from the user, and the output that is generated.

**Fig. 1.**
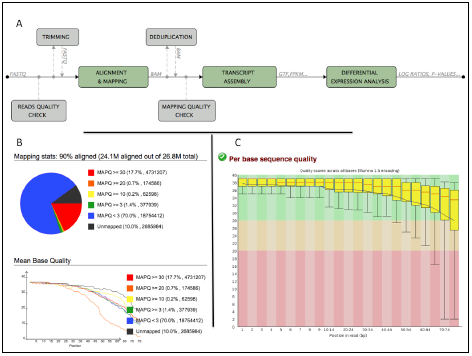
**A.** Concept diagram for RNA-Rocket **B.** SAMStat output showing number of aligned reads. **C.** FastQC profile on base call quality

RNA-Rocket is designed to make users aware of three major concerns: the quality base calls in their sequencing reads, the number of reads aligned from their sample, and accounting for PCR bias. Base call quality can vary depending on the sequencing technology and sample preparation. Poor quality sequencing can impact the certainty with which a read can be mapped to the genome (Li, Ruan, & Durbin, 2008). Although modern aligners endeavor to account for Phred quality scores when performing alignment, it is important for users to consider the quality of their input when performing RNA-Seq analysis since low quality input can lead to low levels of alignment. To promote this awareness, the RNA-Rocket site highlights this quality control step and provides the FastQC tool (Andrews, 2010) for determining the quality profile of sample reads. The site also provides users with access to Sickle (Joshi) and Trimmomatic (Bolger, Lohse, & Usadel, 2014), which can be used to automatically trim off low quality base calls from the ends of sequencing reads. Once alignment is performed the user also has the option to check the quality profile and number of reads mapped using a modified version of the SAMStat tool (Lassmann, Hayashizaki, & Daub, 2011). These three options enable analysis for the researcher to maximize the amount of their sample being used through iterative pre-processing, alignment, and evaluation.

RNA-Rocket also provides a quality control option for removing PCR bias that can occur as a result of library preparation (Aird et al., 2011). Certain sequences may be overrepresented if their composition leads to bias in the amplification process (Benjamini et al.). For paired-end data this option uses Picard tools (“Samtools Picard-Tools,”) to “collapse” multiple read pairs that have identical coordinates for the first and second read into a single representative pair.

In addition to RNA-Seq analysis, RNA-Rocket also supports ChIP-Seq (Chromatin Immunoprecipitation-Sequencing) analysis by providing access to the peak calling program MACS (Zhang et al., 2008). After mapping the reads to the reference genome, a user may use MACS to identify and quantify the ChIP signal enriched genomic regions. The mapped reads and genome coverage can be directly viewed via the BRC genome browsers and the peak calling result can be viewed and downloaded via the RNA-Rocket web interface.

### 2.2 Implementation

The RNA-Rocket site is updated daily with genomes and annotations from each of the contributing BRCs. Thousands of genomes are organized and indexed using Bowtie2 (Langmead & Salzberg, 2012) and SAMtools (Li et al., 2009) to enable alignment and bias correction for abundance estimation respectively. Reference resources are organized by BRC so that results can be streamed and analyzed within the context of the data provider.

The RNA-Rocket project modifies the existing Galaxy code so that Galaxy workflows can be constructed in advance by system administrators and shared to users through a tiered menu system. This menu system, referred to as “Launch Pad”, organizes RNA-Seq processing steps conceptually and gives increasing detail as the user progress towards launching a job i.e. an analysis step. This system also asks the user to populate their project space, known as a ‘history’ in Galaxy terms, with the necessary files before attempting to configure the parameters for their job. This is designed to minimize confusion when attempting to setup an analysis and promote organization for projects involving many files and processing steps.

**Table 1.**
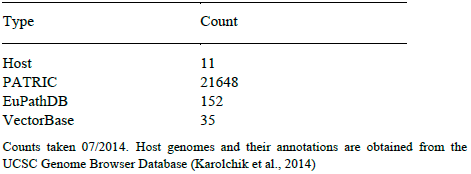
Number of pre-indexed genomes with annotations and streaming support

| Type | Count |
| --- | --- |
| Host | 11 |
| PATRIC | 21648 |
| EuPathDB | 152 |
| VectorBase | 35 |

Counts taken 07/2014. Host genomes and their annotations are obtained from the UCSC Genome Browser Database (Karolchik et al., 2014)

## 3 RESULTS

Brought online in early 2012 RNA-Rocket has steadily increased its user base, the amount of data submitted and generated, and the number of genomes supported and analyzed. As of 07/2014 the site has over 860 registered users, 9Tb of uploaded data and 5Tb of output generated. Multiple research projects have leveraged the free compute resources to process their transcriptomic data (Lew et al., 2013; Stringer et al., 2014) and support continues to expand as requests come in for additional genomes and annotations. The RNA-Rocket site provides a simplified interface targeted for novice users designed to make users aware of the quality control issues and other caveats associated with using high-throughput sequencing data for transcriptomic analysis. Using the workflow system, RNA-Rocket provides pre-formulated solutions for common problems that users encounter. These workflows are easily adapted to new tools and are publicly available for download to enable offline analysis and customization for researchers. By combining experimental concepts with file based requirements the user interface aims to guide life science researchers through the process of RNA-Seq data analysis while making them aware of quality control caveats like base-call quality and unaligned reads.

### 3.1 RNA-Rocket as an educational tool

The RNA-Rocket resource has been, and continues to be, used in numerous scientific workshops throughout the country as a way to familiarize researchers with the steps necessary to process RNA-Seq data. RNA-Rocket provides example data from each of the BRC projects so that users can familiarize themselves with the site using real data. Some of these data are provided through the Driving Biological Project (DBP) initiative, research projects competitively enabled through NIAID’s BRC program designed to drive innovation at the BRC sites based on the needs of the research community. Each dataset is provided as both “before” and “after” project spaces so that users can import the project into their own user space, run their own analysis, and view pre-existing results that demonstrate the analysis capability. These project spaces were also created in conjunction with How-To documents, available at http://rnaseq.pathogenportal.org/page/list_published, that take the user through the step-by-step process of generating output for analysis.

#### 3.1.1 PATRIC educational data

Non-typhoidal *Salmonella* (NTS) are widespread and common human pathogens, causing tens of millions of infections and hundreds of thousands of fatalities annually worldwide, especially among young and elderly victims with co-morbidities. RNA-Rocket makes available transcriptome data generated through a PATRIC Driving Biological Project for *Salmonella typhimurium* 14028. These data seek to characterize the transcriptome of *S. typhimurium* in defined environments. Specifically ten different growth conditions were used: Rich media (LB) logarithmic and stationary phase growth, M9 minimal media logarithmic and stationary phase growth, M9 minimal media containing glucose logarithmic and stationary phase growth. Finally, bacteria at early stationary were compared with and without nitric oxide treatment, which the cell uses to try to kill *Salmonella*. All experiments were performed twice on different days to account for biological and technical variation. Sequencing is paired-end and strand specific using Illumina encoding 1.5 and 1.9.

#### 3.1.2 EuPathDB educational data

EuPathDB has utilized RNA-Rocket as an educational tool to empower bench scientists to perform their own RNA-Seq analysis without the need to master command line scripting. To this end EuPathDB includes a two-section module on RNA-Seq analysis using RNA-Rocket in all of its workshops. In the first section workshop participants learn about RNA-Seq file format and the tools used to align and analyze RNA-Seq data. In addition, they work with a sample dataset available from SRA (Leinonen, Sugawara, & Shumway, 2011), setup TopHat and Cufflinks parameters and kickoff the workflow. In the second section, the results of the workflow are examined including a description of output file formats and conversion between formats. Students then transfer their data to the genome browser in EuPathDB to visualize the data in the context of the genome, annotation and other available functional data. EuPathDB has also developed YouTube tutorials that describe how to use RNA-Rocket, which can be found, along with the workshop module, at the RNA-Rocket site.

To enable users to explore data associated with a eukaryotic pathogen RNA-Rocket provides short read sequences, alignments, and assembled transcripts for *Encephalitozoon cuniculi* (Grisdale, Bowers, Didier, & Fast, 2013), a parasitic organism known to infect humans with compromised immunity due to HIV-infection. Grisdale et al. generated RNA-Seq data for *E. cuniculi* at three time-points: 24 hr, 48 hr, and 72 hr post-infection of rabbit kidney fibroblast, RK13 cells. Short read sequences were obtained from SRA.

#### 3.1.3 VectorBase educational data

The transcriptome of *Aedes aegypti* sampled from neural tissues, and other tissues involved in host-seeking behavior, is profiled in Illumina short read data generated through a VectorBase DBP for this dengue and yellow fever vector. The resulting neurotranscriptome (BioProject accession PRJNA236239 and ID 236239) is a catalogue of all mosquito genes expressed in the central nervous system and head sensory appendages that are thought to be involved in host-seeking behavior; the antenna and rostrum (maxillary palp and proboscis); and the brain, which directs the sensorymotor behavior required for a blood meal. Other tissues included ovaries, which are affected by blood feeding, and the mosquito legs and abdominal tips (ovipositors), known to carry chemosensory sensilla involved in gustatory behaviors such as oviposition. Beyond comparing sex and tissue-specific differences in gene expression, this project sought to examine tissue specific gene expression changes associated with blood-feeding state, and such changes differed between two strains of *A. aegypti* with divergent host preferences. This transcriptome contains 147 SRA experiments; data provided on the RNA-Rocket site contains one library for the brain tissue of females fed with sugar.

### 3.2 BRC features

After results have been computed at the RNA-Rocket site they can be streamed back to the respective BRC depending on the reference organism selected for analysis. This provides users with the ability to process and analyze their RNA-Seq data remotely without having to download potentially large files to their own computer.

#### 3.2.1 PATRIC

PATRIC supports three major data types generated by RNA-Seq analysis at RNA-Rocket through their corresponding file formats: alignments (BAM and bigWig), transcript assembly (GTF), and differential expression analysis data (GeneMatrix file). RNA-Seq data can be streamed directly from RNA-Rocket to the appropriate organism’s genome browser, JBrowse (Skinner, Uzilov, Stein, Mungall, & Holmes, 2009), at PATRIC, allowing researchers to examine these data and compare it with the annotations that are provided for each genome. By streaming the transcript assembly results from RNA-Rocket to PATRIC a user can determine novel genomic features predicted by their RNA-Seq experiment relative to existing PATRIC and NCBI annotations using PATRIC’s ‘Combination Track’ ability. When using RNA-Rocket, novel transcript prediction can be performed in two different ways: assemble transcripts using an existing annotation and assemble transcripts *de novo*. When creating a combination track from transcripts assembled using an existing annotation, a user can employ PATRIC’s ‘subtract features’ option to easily see all genomic features predicted by the RNA-Seq experiment that are not present in the annotation. When using transcripts assembled *de novo* the resulting predictions more closely resemble the operon structure from a given bacteria. In this case a combination track can be created using PATRIC’s ‘subtract coverage’ feature to view all those regions, which were part of a transcribed region of the genome but not in an existing annotation. This allows users to easily encapsulate and view novel predictions made by their RNA-Seq experiment. These combination tracks can then be downloaded as a GFF3, BED, or Sequin table file. As seen in Figure 2, PATRIC’s browser also allows viewing of alignment BAM files, to show alignment details, and BigWig to see an overall expression profile.

**Fig. 2.**
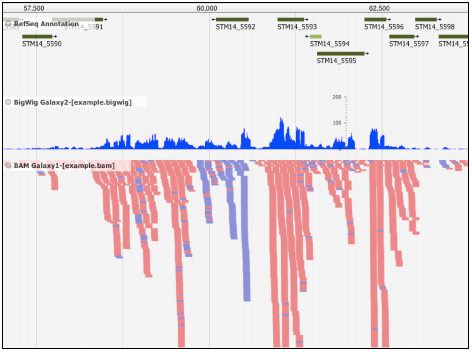
PATRIC view of existing annotation (top) with a BigWig expression profile (middle) and BAM alignment view (bottom)

To support differential gene expression analysis, PATRIC provides a suite of integrated tools to explore, visualize, analyze and compare published and private datasets using a secure workspace. Researchers can upload their own pre-processed transcriptomics datasets, generated by either RNA-Rocket or a microarray experiment to their private workspace and analyze them with data analysis and visualization tools available at PATRIC. Metadata-driven filters facilitate searching for experiments of interest; dynamic filterable gene lists, a heat map, and clustering tools allow researchers to quickly find genes that are up or down regulated in the select experimental conditions or have similar expression patterns across one or more conditions. PATRIC’s pathway summary tool summarizes metabolic pathways related to selected genes. As of July 2014, the PATRIC database included 807 transcriptomics experiments across 72 bacterial genera, each incorporated with manually curated data and metadata, all available for comparison. Filtering on experimental metadata can help researchers locate similar experiments to those in their private data and subsequently look for similarities in up- or down-regulation of genes.

#### 3.2.2 EuPathDB

Results computed on RNA-Rocket for eukaryotic pathogens provide links for streaming to the corresponding EuPathDB page for the organism under analysis. This support extends to all the divisions of EuPathDB including AmoebaDB, CryptoDB, GlardiaDB, PiroplasmaDB, PlasmoDB, ToxoDB, TrichDB, TriTrypDB, and the pan-fungal genome resource FungiDB. EuPathDB supports streaming of RNA-Rocket results using BAM, GTF, and bigWig files through GBrowse. As seen in Figure 3, at EuPathDB a user can view predicted exons and transcripts given by Cufflinks and the expression profile given by BAM and bigWig files computed from short read alignment by TopHat. This enables users to see experimental results alongside EuPathDB annotations. Custom tracks are associated with a users login to make them available between sessions and the snapshot functionality allows users to save combinations of tracks of interest.

**Fig. 3.**
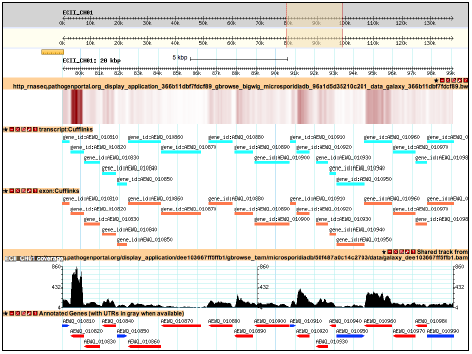
EuPathDB view of RNA-seq data using GBrowse. From top to bottom tracks include nucleotide scale, bigWig density, GTF predicted transcripts and genes, BAM expression view, and EuPathDB annotations

#### 3.2.3 VectorBase

RNA-Rocket supports streaming for 35 different vector genome assemblies from VectorBase. Tracks can be streamed for BigWig files that enable the user to see the expression profile and GTF files which show predicted transcript structure. VectorBase uses the Ensembl browser and supports the ability of users to save custom tracks in association with their user account. For many of the supported organisms, VectorBase provides pre-computed transcriptomic experiments that can be enabled to view alongside a user’s RNA-Seq data. Figure 4 illustrates streaming of the example *A. aegypti* data for brain tissue of females fed with sugar alongside an enabled track that represents a pre-computed result from a female *A. aegypti* brain post-bloodmeal. These data are easily compared to existing VectorBase annotations for protein-coding genes and non-coding RNA to identify potentially undiscovered genomic features.

**Fig. 4.**
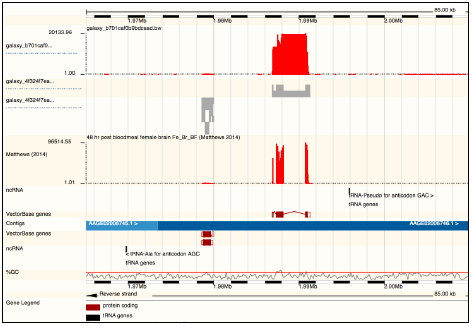
VectorBase view of (top down) bigWig expression profile, GTF RNA-Seq predictions, a pre-computed VectorBase experiment, and VectorBase annotations

## 4 CONCLUSION

RNA-Rocket is a free service that can be used by life-science researchers to process and analyze their RNA-Seq data. The site maintains up-to-date genome and annotation data through NIAD’s Bioinformatic Resource Centers. By leveraging BRC data and the Galaxy system the RNA-Rocket project can provide up-to-date tools and capability despite the rapidly changing landscape of RNA-Seq analysis.

## ACKNOWLEDGEMENTS

The Pathogen Portal team would like to acknowledge the efforts of Alison Yao and Yan Zhang for helping to make this project a reality.

### Funding

This project has been funded with Federal funds from the National Institute of Allergy and Infectious Diseases, National Institutes of Health, Department of Health and Human Services, under Contract No. HHSN272200900040C awarded to BWS Sobral.

